# Recording temporal data onto DNA with minutes resolution

**DOI:** 10.1101/634790

**Authors:** Namita J Bhan, Jonathan Strutz, Joshua Glaser, Reza Kalhor, Edward Boyden, George Church, Konrad Kording, Keith E.J. Tyo

## Abstract

Recording biological signals can be difficult in three-dimensional matrices, such as tissue. We present a DNA polymerase-based strategy that records temporal biosignals locally onto DNA to be read out later, which could obviate the need to extract information from tissue on the fly. We use a template-independent DNA polymerase, terminal deoxynucleotidyl transferase (TdT) that probabilistically adds dNTPs to single-stranded DNA (ssDNA) substrates without a template. We show that *in vitro*, the dNTP-incorporation preference of TdT changes with the presence of Co^2+^, Ca^2+^, Zn^2+^ and temperature. Extracting the signal profile over time is possible by examining the dNTP incorporation preference along the length of synthesized ssDNA strands like a molecular ticker tape. We call this TdT-based untemplated recording of temporal local environmental signals (TURTLES). We show that we can determine the time of Co^2+^ addition to within two minutes over a 60-minute period. Further, TURTLES has the capability to record multiple fluctuations. We can estimate the rise and fall of an input Co^2+^ pulse to within three minutes. TURTLES has at least 200-fold better temporal resolution than all previous DNA-based recording techniques.

## Introduction

Measuring biosignals that span a large range of spatial and temporal scales is critical to understanding complex biological phenomena. In many systems, analytical techniques must probe many cells simultaneously to capture system-level effects, including cells deep in a tissue without disturbing the biological environment^1^. A particularly challenging problem is the measurement of molecules at cellular (or subcellular) spatial resolution and sub-minute temporal resolution in crowded environments. For example, in neuroscience, it is desirable to record neural firing over time across many neurons in brain tissue^2^. Many other recording scenarios are also complex systems, such as in developmental biology^3^ and microbial biofilms^4^, where dynamic waves of signaling molecules determine function. Thus there is a need to study time-dependent biosignals simultaneously in many locations.

To address this need, optical or physical approaches are often employed^5^. However, optical resolution suffers at depth, and physical probes, such as electrodes, can disturb the environment^6^. Furthermore, parallel deployment of multiple probes to simultaneously record data from many cells remains uniquely challenging^1,6,7^. Genetically encoded biorecorders (nanoscale biological devices that record biosignals), specifically those that store information in DNA, represent an attractive alternative. These biorecorders could be delivered to all cells through transgenesis where they are synthesized locally and record in parallel, obviating the challenges of optical and physical methods that must recover the data on the fly across many cells and in deep tissues^1,6,7^.

To date, biorecording strategies that record onto DNA locally and are genetically encodable have been demonstrated with temporal resolution of two hours or more^1,7^. These DNA-editing based techniques primarily rely on nucleases or recombinases, both of which are limited to a temporal resolution on the scale of hours because of the time required for (a) expression of the DNA-modifying enzyme and (b) DNA cleavage and repair to store the data^1^. Moreover, due to the architectures of these recording devices, signals are recorded in a cumulative (or on/off) fashion (Fig. 1A). Cumulative signals can determine the amount of a signal a biorecorder was exposed to, but not the specific times of exposure. It is important to deliver biosignal measurements with higher temporal resolution and higher information content.

**Figure 1:**
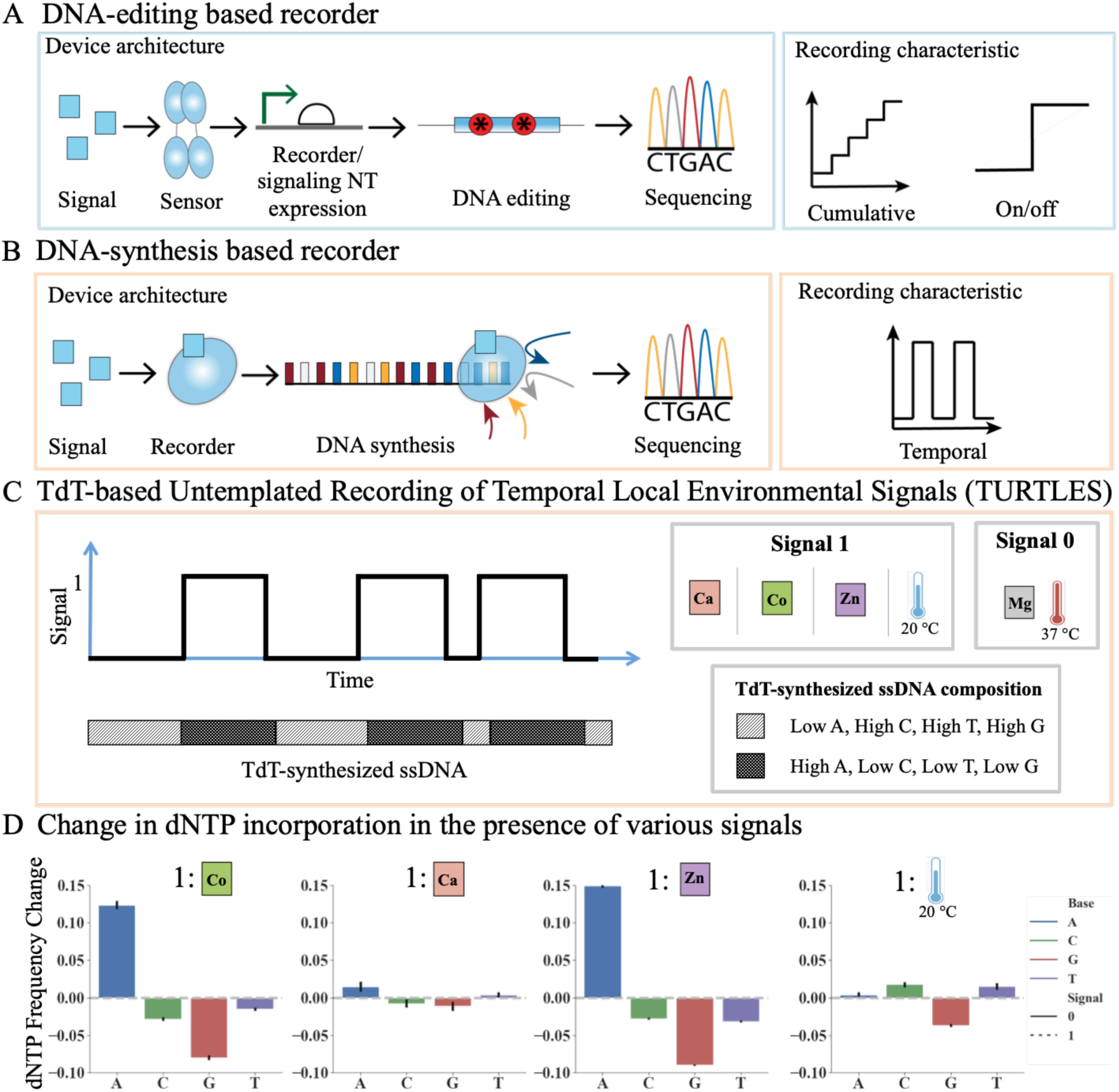
Device architecture of our TdT-based recording system (TURTLES) and its response to various environmental signals. (A) General device architecture and recording characteristic of DNA-editing based signal recorders. (B) General device architecture and recording characteristic of DNA synthesis based recorder. (C) General description of TdT-based untemplated recording of temporal local environmental signal (TURTLES). A time-varying input signal results in synthesis of ssDNA by TdT with varying dNTP compositions (shown as diagonal stripes for signal 0 and crisscross for signal 1). The various signals tested are shown as signal 1 and the background condition shown as signal 0. (D) Change in frequency of dATP, dCTP, dGTP and dTTP incorporation by TdT in the presence or absence of various signals. Signal 0 is always 10 mM Mg^2+^ at 37 °C for 1 hour. Signal 1 was, going from left to right: (1) 10 mM Mg^2+^ + 0.25 mM Co^2+^ at 37 °C for 1 hour; (2) 10 mM Mg^2+^ + 1 mM Ca^2+^ at 37 °C for 1 hour; (3) 10 mM Mg + 20 μM Zn^2+^ at 37 °C for 1 hour; and (4) 10 mM Mg^2+^ at 20 °C for 1 hour. Error bars show two standard deviations of the mean. Statistical significance was assessed after first transforming the data into Aitchison space which makes each dNTP frequency change statistically independent of the others (fig. S2).

Terminal deoxynucleotidyl transferases (TdTs) belong to a unique class of DNA polymerases (DNAp) that synthesize single stranded DNA (ssDNA) in template-independent fashion^8,9^. TdTs incorporate dNTPs probabilistically to the 3’ termini of ssDNA substrates according to an inherent dNTP incorporation preference^10^. As we show, this dNTP preference is affected by changes in the TdT reaction environment. When the dNTP incorporation preference is altered, then information about the environment could be recorded in each incorporated dNTP. We thus prototype a DNA-synthesis based biorecorder for achieving the spatiotemporal resolution that eludes the current DNA-editing based biorecorders.

We present TdT-based Untemplated Recording of Temporal Local Environmental Signals (TURTLES). We demonstrate that TURTLES can achieve minutes temporal resolution (a 200-fold improvement over existing DNA recorders) and outputs a truly temporal (rather than cumulative) signal. We show that changes in divalent cation concentrations (Ca^2+^, Co^2+^, and Zn^2+^) and temperature alter dNTP incorporation preferences of TdT and that concentrations and temperatures can be recovered by analyzing the ssDNA synthesized by TdT. We show that temporal information can be obtained by using estimates of dNTP incorporation rates, allowing us to map specific parts of a DNA strand to moments in time of the recording experiment. Using this approach, we can record temperature and divalent cation dynamics with a few minutes frequency. The current *in vitro* study demonstrates the potential for TdTs as DNA-based biorecorders with high temporal resolution.

## Results

### TdT can detect environmental signals *in vitro* via changes in dNTP incorporation preference

For TdT, the kinetics of incorporation for specific nucleotides is affected by the cations present in the reaction environment^9^. For example, previous studies^8,11,12^ and our experiments show that when only one nucleotide is present, TdT incorporation rates of pyrimidines, dCTP and dTTP, increase in the presence of Co^2+^ (fig. S1).

We sought to examine if Co^2+^-dependent changes in kinetics also occurred in the presence of all four nucleotides, dATP, dCTP, dGTP, and dTTP (hereon referred to as A, C, G, and T). ssDNA substrate extended by TdT in the presence of Mg^2+^ only and with 0.25 mM CoCl_2_ added were determined by single molecule sequencing. Upon Co^2+^ addition, A incorporation increased by 13%, while G decreased by 10% and T and C decreased by 3 and 2 percent respectively (these values do not sum to 0% due to rounding error) (Fig. 1D and fig. S2). This shift in dNTP incorporation preference could be used to determine if Co^2+^ was present or not during ssDNA synthesis.

Next, we were interested in understanding how many biologically relevant signals could be recorded by TURTLES. We examined Ca^2+^, Zn^2+^, and temperature. Ca^2+^ is a proxy for neural firing^13^, Zn^2+^ is an important signal in development and differentiation of cells^14^, and temperature is relevant in many situations.

Different environmental signals had differences both in the particular dNTP affected and the magnitude of the dNTP incorporation preference change. For 20 μM Zn^2+^, we saw a 15% increase in a preference for A, 8% decrease in a preference for G, 4% decrease in a preference for T, and 3% decrease in a preference for C (Fig. 1D and fig. S2). dNTP incorporation preference upon 1 mM Ca^2+^ addition changed more modestly. The change was 1.4% increase in A, 1.7% decrease in G, 1.0% increase in T and 0.5% decrease in incorporation of C (Figure 1D and fig. S2). Finally, we changed the reaction temperature from the preferred 37 °C to 20 °C and saw a 3% increase in A, 3.5% decrease in G, 1.0 % increase in T and 0.5% decrease in incorporation of C (Figure 1D and fig. S2). The addition of cations as well as temperature change altered the dNTP incorporation rates and lengths of ssDNA strands synthesized (fig. S3-8). Thus, we were able to characterize the effect of multiple biologically relevant signals on TdT activity. For further analysis of TURTLES we chose to focus on Co^2+^ as the candidate cationic signal.

### Recording a single step change in Co^2+^ concentration onto DNA with minutes resolution *in vitro*

Having quantified the distinct change in dNTP incorporation preference upon Co^2+^ addition, we next examined if we could identify the time at which Co^2+^ was added to a TdT-catalyzed ssDNA synthesis reaction based on the change in sequence of the synthesized ssDNA strands (Fig. 2, A and B). During a 60 min extension reaction, we created input unit step functions at 10, 20, and 45 minutes by adding 0.25 mM Co^2+^ at those times (we will refer to this as a 0➔1 signal where ‘0’ is without Co^2+^ and ‘1’ is with 0.25 mM Co^2+^). In order to test the resolving power of TURTLES, we sought to infer these times from the DNA readout. For each reaction, we analyzed approximately 500,000 DNA strands by single molecule sequencing and calculated the dNTP incorporation frequencies over all reads. By plotting the change in dNTP incorporation frequency along the extended strands after normalizing each sequence by its own length, we showed that later addition of Co^2+^ resulted in changes farther down the extended strand (Fig. 2C). We then calculated the average location across all the sequences for a given condition at which half the 1 control (Mg^2+^+Co^2+^) signal was reached. To translate this location into a particular time in the experiment, we assumed constant rate of dNTP addition (fig. S9) and derived an equation that adjusted for the change in rate of DNA synthesis between the 0 and 1 controls (Equation 5, Materials and Methods). Using this information, we could estimate the Co^2+^ additions to be at 9.9, 21.5 and 46.6 minutes (Fig. 2D). We were also able to estimate the time within 7 minutes of the unit input step function for the reverse; a change in signal (Co^2+^ concentration) from 1 to 0 (fig. S10). We thus show that TURTLES has excellent temporal precision, approximately 200-fold higher than any other currently utilized biorecorders.

**Figure 2:**
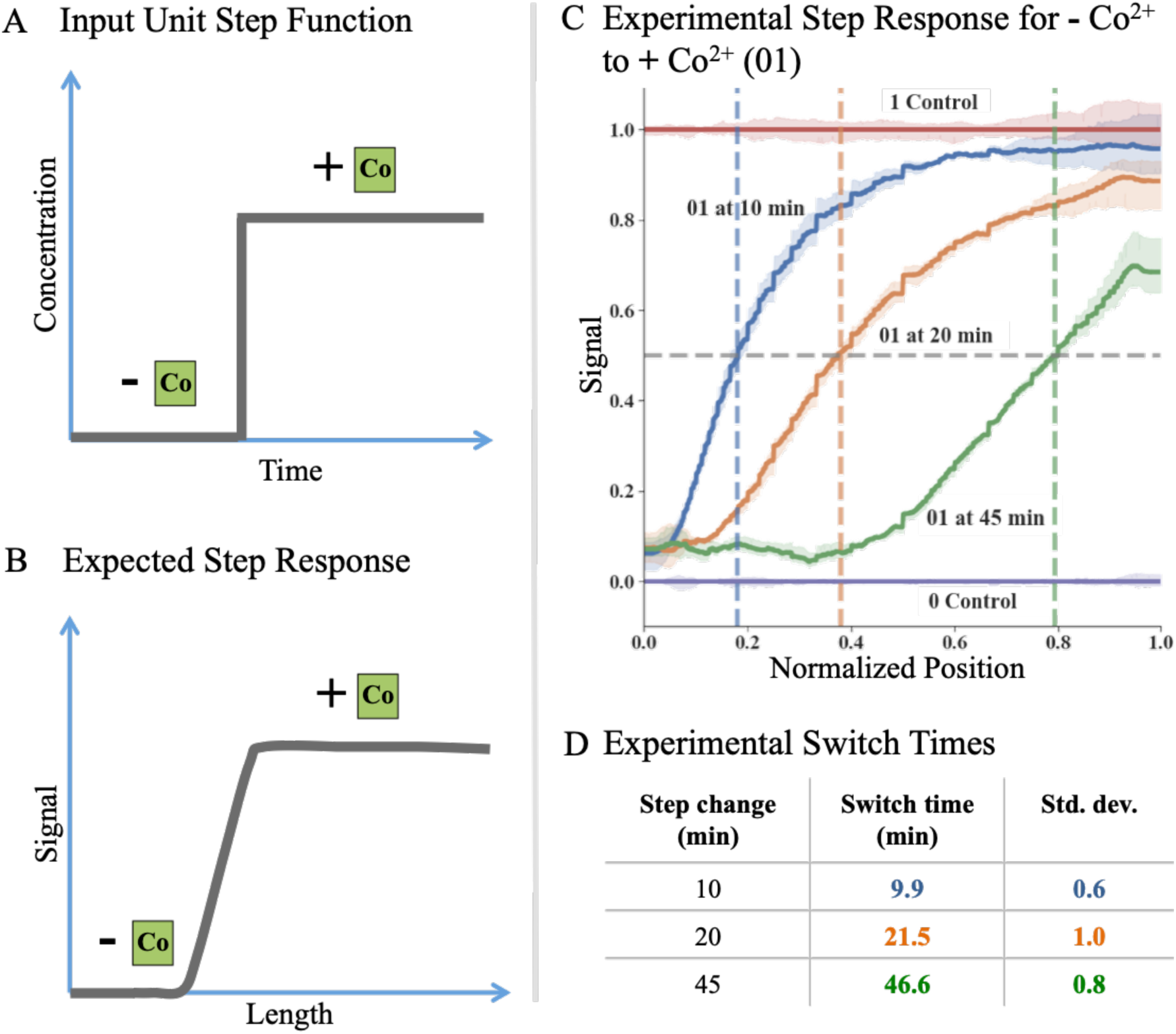
Recording a single step change in Co^2+^ concentration onto ssDNA with minutes resolution *in vitro.* (A) Representative input unit step function used in our experiments by changing concentration of Co^2+^ from 0 mM to 0.25 mM during a TdT-based DNA synthesis reaction while keeping Mg^2+^ concentration and reaction temperature constant. (B) Expected step response of the TdT-based DNA recording system for the 0➔1 input unit step function. (C) Experimental data for various input unit step functions each with 0.25 mM Co^2+^. Signal is calculated based on differences in dNTP preference. This plot shows there is a difference in the preference of dNTP incorporated by TdT in the Mg^2+^ (purple) and Mg^2+^+Co^2+^ (red) control conditions (where the signal (Co^2+^) is not added or removed throughout the extension reaction). The plot further shows the changes from 0➔1 for Co^2+^ added at 10 minutes (blue), Co^2+^ added at 20 minutes (orange), and Co^2+^ added at 45 minutes (green). Total extension time for each of these experiments was 60 minutes. (D) Table showing the actual switch time as well as the mean inferred switch time along with each mean’s standard deviation (mean calculated across 3 biological replicates).

While we were able to accurately estimate the times of Co^2+^ addition (0➔1) and removal (1➔0), in many applications, simultaneously synthesizing ~500,000 strands of DNA will be infeasible. To determine the number of strands needed for reasonable statistical certainty, we randomly sampled smaller groups of strands from the experiment and evaluated our ability to predict when Co^2+^ was added (fig. S11 and S12). With only ~6,000 strands, we still estimated the time of Co^2+^ additions to be at 9.7, 23.2 and 44.7 minutes (table S1). Thus, even with a limited number of strands, high temporal precision recording is feasible.

### Recording multiple fluctuations in Co^2+^ concentration onto DNA with minutes resolution *in vitro*

As mentioned, an advantage of this approach is that it can record the time of multiple fluctuations. This is in contrast to any of the other DNA-based recorders, which rely on an accumulation of signal (i.e. accumulation of mutations). Accumulation can tell what fraction of the time a signal was present over a period of time, but not how the signal was distributed throughout the time period of recording. The ability to know when fluctuations occur would allow new levels of insight into different biological systems.

We used TURTLES to record a 0➔1➔0 signal, where ‘0’ is without Co^2+^ and ‘1’ is with 0.25 mM Co^2+^ (Fig. 3A). The signal was 0 for the first 20 minutes, 1 for the next 20 minutes, and 0 for the last 20 minutes of the extension reaction (Fig. 3A). We took the sequencing data obtained from the experiment and calculated the signal (Fig. 3B). Because multiple step changes were present, we used an algorithm previously developed by Glaser et al.^15^ (see Materials and Methods for details) to estimate the true value of the signal at all times (every 0.1 min). Impressively, the signal reconstruction clearly resembles the true 0➔1➔0 signal, with transitions between the 0 and 1 signals occurring at 23.2 and 40.7 minutes (Fig. 3C). The possible reason for the modest increase in variability is discussed in sup. text 5 and fig. S13. Finally, using *in silico* simulations based on the experimental parameters of TdT, it is clear that a TdT-based recording system can accurately record more than 3 pulses and pulses of much shorter duration than 20 minutes (fig. S14). Overall, this demonstrates the capability of TURTLES to record multiple temporal fluctuations.

**Figure 3:**
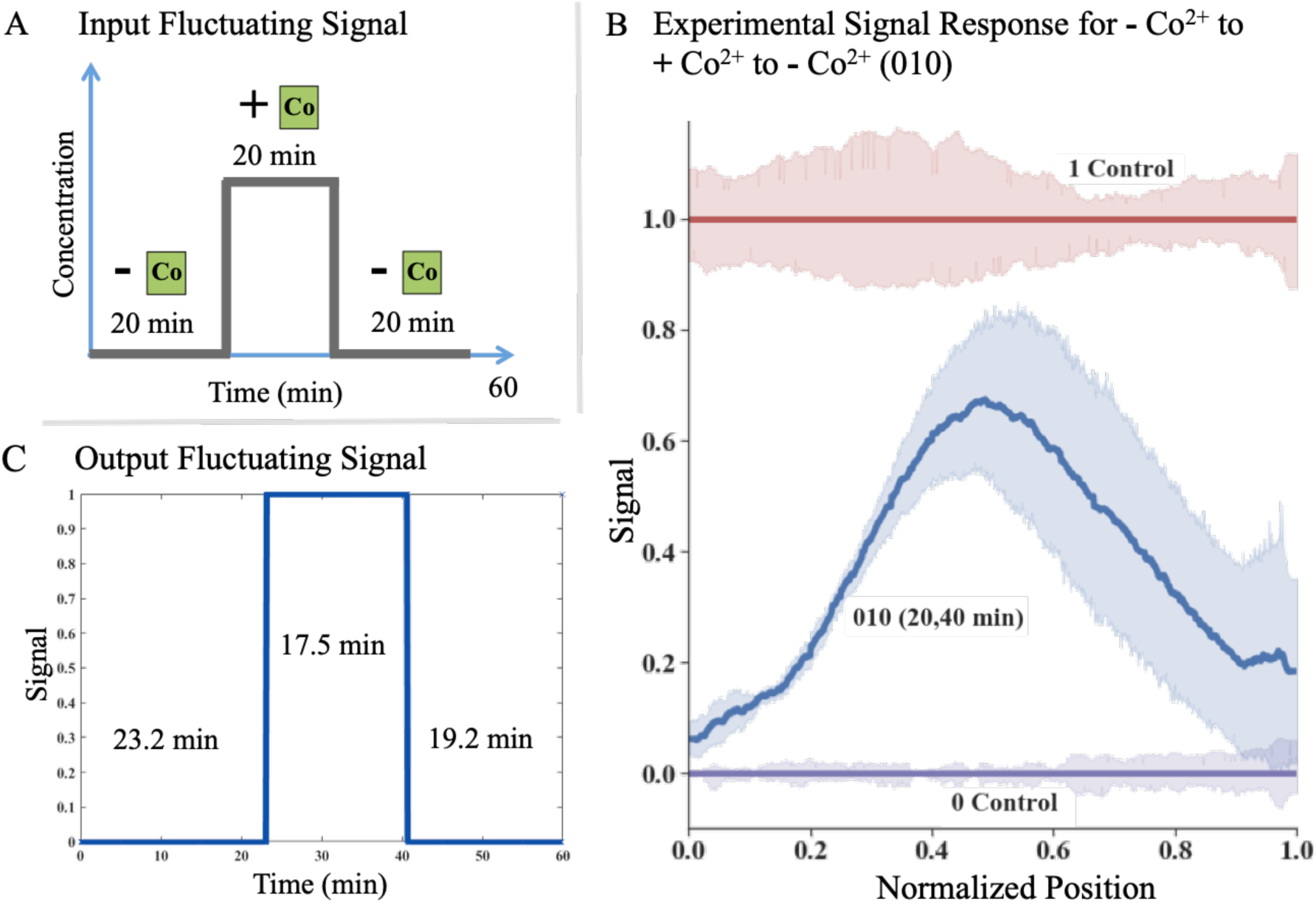
Recording multiple fluctuations of signal onto DNA. (A) Representative fluctuating input signal used in our experiments by changing concentration of Co^2+^ from 0 mM to 0.25 mM and back to 0 mM during a TdT-based ssDNA synthesis reaction while keeping Mg^2+^ concentration and reaction temperature constant. (B) Experimental data for fluctuating input signal of 0 mM Co^2+^➔0.25 mM Co^2+^➔0 mM Co^2+^ (010). Signal is calculated based on differences in dNTP preference. This plot shows there is a difference in the preference of dNTP incorporated by TdT in the Mg^2+^ (purple) and Mg^2+^+Co^2+^ (red) control conditions (where the signal (Co^2+^) is not added or removed throughout the extension reaction). The plot further shows the changes from 0➔1➔0 for Co^2+^ added at 20 minutes and removed at 40 minutes (blue). Total extension time for these experiments was 60 minutes. (C) Output fluctuating signal. Using the algorithm detailed in Glaser et al.^15^, the signal was deconvoluted into a binary response, with predicted switch times of 23.2 minutes and 40.7 minutes (actual: 20 minutes and 40 minutes). Signal predictions were made every 0.1 minutes and lines were added at the times of rise and fall of pulse for visualization.

## Discussion

Our results show that TURTLES can record temporal changes in divalent cationic concentrations and temperature onto DNA at minutes timescale resolution *in vitro.* The methodology we present here is two orders-of-magnitude faster than any of the currently utilized DNA-based environmental signal recording techniques. This enhancement in temporal resolution is because our biorecorder does not rely on temporal expression of DNA-modifying enzymes or DNA repair processes and is simply limited by (a) the incorporation rate of TdT, which is 1 dNTP per second under optimal conditions and (b) the magnitude of the dNTP incorporation preference change. Because this recording system can fully switch from one state and back to the original, the information recording is truly temporal instead of cumulative, unlike nuclease/recombinase based recording techniques^1^.

As with all DNA-based recording schemes, TURTLES can be encoded genetically, and be employed to record and store information locally in DNA with single cell resolution in tissues, where recovering information in real time is challenging via optical or electronic approaches. Adding a unique barcode to each cell being studied can simplify recovery of spatial resolution^16^. Moreover, based on our previous calculations of the metabolic burden on a cell expressing such a *de novo* DNA recording system^6^; given its current signal recording capability and resolution would make recording 10s of temporal events in a single experiment metabolically feasible.

Several opportunities exist to improve the biorecording performance of TdTs. In this study we utilized simple methods for transforming sequence data into temporal information, but more sophisticated models could extract higher-order effects from the sequence data to improve prediction. Additionally, protein engineering could improve the magnitude of dNTP incorporation preference change and reduce the number of simultaneous DNA strands needed for recording. These modifications will be important in cases, like Ca^2+^, where the magnitude of dNTP incorporation preference change is relatively small. As well, protein engineering could improve temporal resolution by enhancing the ssDNA synthesis rate of TdT.

Several attempts are being made to reduce the cost of DNA synthesis associated with phosphoramidite chemistry^17,10^. *In vitro* TdT-based recorders could allow the storage of arbitrary digital information into DNA by controlling the environment to record ‘1s’ and ‘0s.’ For example, a low temperature could be ‘0’ and a high temperature could be ‘1’. Because of the probabilistic nature of TdTs, we could not achieve single base resolution, however 1 bit per 10 bases appears to be currently possible. As such TUTRLES could provide a cheaper more environmentally friendly option for DNA data storage.

In the current study, we have only evaluated TdT *in vitro.* Several hurdles will need to be overcome before it can be deployed *in vivo.* While creating many 3’hydroxy ssDNA substrates for TdT should be feasible *in vivo*^18^, ensuring the extended ssDNA is not acted on by any native nucleases may be challenging. Likewise, other environmental signals that affect dNTP incorporation preference may be fluctuating simultaneously during the experiment creating noise and uncertainty in the reading. Mitigating noise may require additional protein engineering to enhance the specificity of the dNTP incorporation changes to only one or very few environmental signals.

TdT-based DNA recording is a promising technology for interrogating biological systems, such as the brain, where high temporal and spatial resolution is needed. In such systems measurement across many cells are required, and the depth of tissue prevents extracting measurements on the fly from using physical and optical methods. Thus TURTLES provides many exciting opportunities for recording complex biological processes that are infeasible today.

## Acknowledgements

The authors would like to acknowledge Marija Milisavljevic for help with some experiments, and Bradley Biggs and Alec Castinado for helpful discussions and comments on the manuscript. This research was supported in part through the computational resources and staff contributions provided for the Quest high performance computing facility at Northwestern University, which is jointly supported by the Office of the Provost, the Office for Research, and Northwestern University Information Technology. All the next generation sequencing was done with the help of the Next Generation Sequencing Core facility at University of Illinois at Chicago. This work was funded by the National Institutes of Health grants R01MH103910 (to KEJT, KPK, ESB and GMC), and UF1NS107697 (to KEJT, KPK, ESB) and National Institutes of Health Training Grant (T32GM008449) through Northwestern University’s Biotechnology Training Program (to JS). NB and KEJT developed the concept. NB, JS, JG, RK, EB, GC, KK, and KT designed experiments. NB performed all experiments. NB, JS, and JG did primary analysis on DNA sequencing data. All authors contributed to data analysis and preparation of the manuscript.

