## Supplemental Information for "Recording temporal data onto DNA with minutes resolution"

This file includes:

**Materials and Methods**

**Supplementary Text**

**Figs. S1 to S14**

**Table S1**

Table of Contents

**Materials and Methods	3**

Enzymes and ssDNA substrate	3

Extension reaction set-up for reactions analyzed by Next Generation Sequencing (NGS) 3

Extension reaction for calculating affect of Co^2+^, Ca^2+^, Zn^2+^, and temperature on overall dNTP preference of TdT	3

Extension reactions for 0🡪1 set-up	3

Extension reactions for 0🡪1🡪0 set-up	3

ssDNA wash for replacing buffers for 0🡪1🡪0 reactions	4

Illumina library preparation and sequencing	4

NGS data preprocessing	5

Timepoint prediction for 0🡪1 single step change experiments	5

Timepoint prediction for 0🡪1🡪0 multiple fluctuations experiment	7

*In silico* simulations of experiments with multiple pulses	7

**Supplementary Text	8**

1. Extension reaction with individual dNTPs for testing effect of Co^2+^	8

2. Extension reactions for 1🡪0 set-up	8

3. Derivation of Equation 5	8

4. Extensions reaction set-up for calculating rate of dNTP addition	9

5. 0🡪1 at 10 minutes test set-up for checking ssDNA clean-up kit bias	9

**Supplementary Figures	10**

Figure S1: Testing change in individual dNTP preference upon Co^2+^ addition	10

Figure S2: Change in Aitchison distance for dATP, dCTP, dGTP and dTTP incorporation by TdT in the presence or absence of various signals	11

Figure S3: Length distribution of extensions upon addition of Co^2+^ based on NGS data	12

Figure S4: Length distribution of extensions upon addition of Zn^2+^ as seen on ssDNA gel	13

Figure S5: Length distribution of extensions upon addition of Zn^2+^ based on NGS data	14

Figure S6: Length distribution of extensions upon addition of Ca^2+^ as seen on ssDNA gel	15

Figure S7: Length distribution of extensions upon addition of Ca^2+^ based on NGS data	16

Figure S8: Length distribution of extensions upon using temperature as a signal based on NGS data	17

Figure S9: Anomalous dNTP composition initially found at the end of reads and rate of reaction measured for extensions with only Mg^2+^ present	18

Figure S10: Recording a single 1🡪0 step change in Co^2+^ concentration onto ssDNA *in vitro*	19

Figure S11: Mean % error in time prediction for 0🡪1 (Mg^2+^ to Mg^2+^+Co^2+^) data when different proportions of experimental data are used for time prediction (data is randomly sampled)	20

Figure S12: Plots showing 0🡪1 data when different percentages of experimental data were randomly sampled	21

Figure S13: dNTP bias & variablity introduced by ssDNA wash columns	22

Figure S14: *In silico* characterization of shortest resolvable pulses and highest number of consecutive pulses that could be resolved using TURTLES	23

**Supplementary Table	24**

Table S1: Table showing time predictions obtained for 0🡪1 data when different percentages of experimental data were randomly sampled	24

Materials and Methods

**Enzymes and ssDNA substrate:**

Terminal deoxynucleotidyl polymerase, T4 RNA ligase I, Phusion High-Fidelity PCR Master Mix with HF Buffer were purchased through New England Biolabs (NEB). ssDNA substrates used for extension reactions were ordered from Integrated DNA Technologies (IDT) with standard desalting. dNTPs were obtained from Bioline.

**Extension reaction set-up for reactions analyzed by Next Generation Sequencing (NGS)**

**Extension reaction for calculating affect of Co^2+^, Ca^2+^, Zn^2+^, and temperature on overall dNTP preference of TdT:**

Each extension reaction consisted of a final concentration of 10 µM ssDNA substrate (CS1: 5’ACACTGACGACATGGTTCTACA3’), 1mM dNTP mix (each dNTP at 1 mM final concentration), 1.4x NEB TdT reaction buffer, and 10 units of TdT to a final volume of 50 μL. When testing the effect of cations, CoCl_2_ was added at a final concentration of 0.25 mM, CaCl_2_ at 1 mM, or Zn(Ac)_2_ at 20 μM. For CoCl_2_ and Zn(Ac)_2_ reactions were run in triplicates. For CaCl_2_, since the change in dNTP preference was relatively small, seven biological replicates were run, and a second set of seven replicates were run on a separate day. It is important to note that reaction initiation was done by adding TdT to the ssDNA substrate mix (ssDNA substrate mix consisted of the ssDNA substrate, dNTPs and the cation). Prior to reaction initiation, the ssDNA substrate mix and TdT were stored in separate PCR strip tubes at 0 °C (on ice). The reaction was run for 1 hour at 37 °C in a Bio-Rad PCR block. When testing the effect of temperature, the same reaction mix was run on a Bio-Rad PCR block set at tested temperatures in biological triplicates for 1 hour. Reaction was stopped by freezing at -20 °C. For initial testing, 2 μL of the reaction was mixed with 12 μL of TBE-Urea (Bio-Rad) loading dye and boiled for 10 minutes at 100 °C. All of the diluted extension reaction was then loaded onto 30 μL, 10 well 10% TBE-Urea Gel (Bio-Rad) and run for 40 minutes at 200 V. Immediately after the run was over, the gel was stained with Sybr Gold for 15 minutes and imaged on an ImageQuant BioRad.

**Extension reactions for 0🡪1 set-up:**

Mg^2+^ only for 1 hour (signal 0) and Mg^2+^+Co^2+^ for 1 hour (signal 1) were set up as regular extension reaction mentioned above. The 0🡪1 reactions where the signal changed from 0 to 1 at various times during the 1 hour extension were run starting at a total volume of 45 μL with Mg^2+^ only. 5 μL 2.5 mM CoCl_2_ was added at the time we wanted the signal to change from 0 to 1. Reactions were all run for a total of 1 hour in triplicates. Fresh signal 0 and signal 1 controls were run with each set-up.

**Extension reactions for 0🡪1🡪0 set-up:**

Mg^2+^ only for 1 hour (signal 0) and Mg^2+^+Co^2+^ for 1 hour (signal 1) were set up as regular extension reaction mentioned above. The 0🡪1🡪0 reactions where the signal changed from 0 to 1 at 20 minutes and back to 0 at 40 minutes were run starting at a total volume of 45 μL with Mg^2+^ only. 5 μL 2.5 mM CoCl_2_ was added at the time we wanted the signal to change from 0🡪1. For changing the signal from 1🡪0, since the ssDNA was suspended in reaction buffer for these set-ups, we used a ssDNA clean up kit (methods mentioned below) to remove the reaction buffer, TdT, cation and dNTPs from each reaction. All of the ssDNA collected from the ssDNA clean up kit (20 μL) was then prepared for the last part of the extension reaction. Collected ssDNA was mixed with a dNTP mix at a final concentration of 1 mM (each dNTP at 1 mM final concentration), 1.4x TdT reaction buffer and 10 units of TdT to a final volume of 50 μL. All reactions were always initiated by adding TdT in the end. Signal 0 and signal 1 controls were run for 1 hour for each set-up in triplicates and also put through the ssDNA wash step at 40 minutes. Six replicates were run for 0🡪1🡪0 reactions.

**ssDNA wash for replacing buffers for 0🡪1🡪0 reactions:**

For changing cation concentration from 1 to 0 we utilized the ssDNA clean-up kit (ssDNA/RNA clean/concentrator D7010) from Zymo Research such that all the extended ssDNA synthesized in the initial part of the experiment was retained on the column and the TdT, reaction buffer, cation and dNTPs were washed away. Each 50 μL extension reaction was individually loaded into a separate column. Protocol was followed as mentioned in the kit. ssDNA was eluted into 20 μL ddH_2_O. We noticed in initial tests that after using the ssDNA clean-up kit, there was little to no TdT-based extension in some replicates (data not included). We presume this is due some ethanol getting carried forward into the eluted ssDNA. Thus we extended the dry spin time based on suggestion from Zymo Research to 4 minutes. We also utilized two other ways to evaporate any remaining ethanol after the column dry spin step based on protocol mentioned in Cold Spring Harbor Protocols^1^. We either kept the columns open in a biohood for 15 minutes to allow for evaporation, or after elution of ssDNA we kept the 1.5 mL eppendorf tubes containing the eluted ssDNA open at 45 °C for 3 minutes. Both methods gave better ethanol removal than just dry spin, and they were tried in triplicates and averaged and plotted for the time prediction analysis (Fig. 3C).

**Illumina library preparation and sequencing:**

Our sample preparation pipeline for NGS was adapted from a previous protocol^2,3^. After extension reaction, 2 μL of the product was utilized for a ligation reaction. 22 bp universal tag, common sequence 2 (CS2) of the Fluidigm Access Array Barcode Library for Illumina Sequencers (Fluidigm), synthesized as ssDNA with a 5’ phosphate modification and PAGE purified (Integrated DNA Technologies), was blunt-end ligated to the 3’ end of extended products using T4 RNA ligase. Ligation reactions were carried out in 20 μL volumes and consisted of 2 μL of extension reaction, 10 μL of 50% PEG 8000, 1 mM ATP, 1 μM CS1 ssDNA, 1X T4 RNA Ligase Reaction Buffer (NEB), and 10 units of T4 RNA Ligase 1 (NEB). Ligation reactions were incubated at 25 °C for 16 hours. Ligated products were stored at −20 °C until PCR that was carried out on the same day. Ligation products were never stored at -20 °C for more than 24 hours.

PCR was performed with barcoded primer sets from the Access Array Barcode Library for Illumina Sequencers (Fluidigm) to label extension products from up to 96 individual reactions. Each PCR primer set contained a unique barcode in the reverse primer. From 5’-3’ the forward PCR primer (PE1 CS1) contained a 25-base paired-end Illumina adapter 1 sequence followed by CS1. The binding target of the forward PCR primer was the reverse complement of the CS1 tag that was used as the starting DNA substrate. From 5’-3’ the reverse PCR primer (PE2 BC CS2) consisted of a 24-base paired-end Illumina adapter 2 sequence (PE2), a 10-base Fluidigm barcode (BC), and the reverse complement of CS2. CS2 DNA that had been ligated onto the 3’ end of extended products served as the reverse PCR primer-binding site. Each PCR reaction consisted of 2 μL of ligation product, 1X Phusion High-Fidelity PCR Master Mix with HF Buffer (NEB), and 400 nM forward and reverse Fluidigm PCR primers in a 20 μL reaction volume. Products were initially denatured for 30 s at 98 °C, followed by 20 cycles of 10 s at 98 °C (denaturation), 30 s at 60 °C (annealing), and 30 s at 72 °C (extension). Final extensions were performed at 72 °C for 10 min. Amplified products were stored at −20 °C until clean up and pooling. QC for individual sequencing libraries was performed as follows. 2 μL of each library was pooled into a QC pool and the size and approximate concentration was determined using Agilent 4200 Tapestation. Pool concentration was further determined using Qubit and qPCR methods. Sequencing was performed on an Illumina MiniSeq Mid Output flow cell and sequencing was initiated using custom sequencing primers targeting the CS1 and CS2 conserved sites in the library linkers. Additionally phiX control library was spiked into the run at 15-20% to increase diversity of the library clustering across the flow cell. After demultiplexing, the percent seen for each sample was used to calculate a new volume to pool for a final sequencing run with evenly balanced indexing across all samples. This pool was sequenced with metrics identical to the QC pool. Library preparation and sequencing were performed at the University of Illinois at Chicago Sequencing Core (UICSQC).

**NGS Data Preprocessing:**

For each sample, the NGS reads were first trimmed and filtered using cutadapt (v1.16). Only NGS read pairs with both Illumina Common Sequence adapters, CS1 and CS2, were kept. Of these, CS2 was trimmed off each R1 sequence and CS1 was trimmed off each R2 sequence. Cutadapt parameters were set as following: a minimum quality cutoff (-q) of 30, a maximum error rate (-e) of 0.05, a minimum overlap (-O) of 10, and a minimum extension length (-m) of 1. The minimum overlap was set to be higher than the default value of 3 because extended sequences in this case are random, and we did not want to filter out sequences where the final 1-10 bases just happen to look like the first 10 bases of CS2 (the read must still contain a full CS2 sequence for it to be kept and subsequently trimmed, however). The 3’ (-a) adapter trimmed from the R1 reads was 5’AGACCAAGTCTCTGCTACCGTA3’ (CS2 reverse complement), and the 5’ (-A) adapter trimmed from the R2 reads was 5’TGTAGAACCATGTCGTCAGTGT3’ (CS1 reverse complement). FastQC was used to quickly inspect the output trimmed .fastq files before downstream analysis. See *filter_and_trim_TdT****.****sh* at <https://github.com/tyo-nu/turtles> for an example preprocessing script. All runs were trimmed using this script. All initial preprocessing was done on Quest, Northwestern University’s high-performance computing facility, using a node running Red Hat Enterprise Linux Server release 7.5 (Maipo) with 4 cores and 4 GB of RAM, although only 1 core was used. Preprocessing took between 5 and 30 minutes depending on the number of conditions, replicates, and reads per replicate in a given run.

Finally, for each analysis, we did further preprocessing locally. We cut off bases that were still present in the reads but not added during the experiment. Degenerate bases (if any) that are part of the 5’ ssDNA substrate (at its 3’ end before the extension) were removed from the beginning of each sequence. Then, we cut off 5.8 bases off the end of every sequence because we found that, on average, 5.8 bases were being added after the extension reaction during the 16-hour ligation step (fig. S9). Because 5.8 is not an integer value, we cut 5 bases off of 80% of the sequences and 6 bases off of 20% of the sequences. We also filtered out sequences with length less than 6 bases.

**Timepoint prediction for 0🡪1 single step change experiments:**

All further analysis was done in python using Jupyter Notebooks. All Jupyter Notebooks used for this publication can be found at [https://github.com/tyo-nu/turtles](https://github.com/tyo-nu/nextgen4b). The following algorithm was applied in order to (1) read and normalize each sequence by its own length, (2) calculate a distance metric using the relative dATP, dCTP, dGTP, and dTTP percent incorporation changes between each condition and the 0 control, and (3) transform distances for all conditions into 0 🡪 1 space based on the 0 and 1 control distance values.

We first normalize each sequence by length, such that all bases in each sequence are counted across 1000 bins. For example, for a sequence of length 10, the first base would get counted in the first 100 bins, the next base in bins 100-200, and so on.

We then calculate base composition, $X_{ij}$, in the sequence for condition, $i$, at each bin with position, $j$, using the formula for a closure (equation 1). Note that $i$ is unique for each (condition, replicate) pair if multiple replicates are present for a given experimental condition.

$$X_{ij}=\left[ \frac{n_{ijA}}{\sum_{k\in N} n_{ijk}},\frac{n_{ijC}}{\sum_{k\in N} n_{ijk}},\frac{n_{ijG}}{\sum_{k\in N} n_{ijk}},\frac{n_{ijT}}{\sum_{k\in N} n_{ijk}} \right] (1)$$

Here, $n_{ijk}$ is the total count of dATP, dCTP, dGTP, or dTTP depending on the value of $k$ ($k\in N=\left\{ A,C,G,T \right\}$) across all sequences for condition, $i$, at bin, $j$.

To calculate distance between two compositions at a given bin location (e.g. between the 0 and 1 controls at every bin), we have to first transform the compositional data. We cannot simply take the L2 norm difference of each compositional element because the elements of a composition violate the principle of normality due to the total sum rule (all elements add up to 100%). Thus, the data is first transformed by using the center log-ratio (clr) transformation which maps this 4-component composition from a 3-dimensional space to a 4-dimensional space. We then take the L2 norm of these transformed normal elements. This distance metric is known as the Aitchison Distance, which is used here to calculate the base composition distance, $d_{j}(0,i)$, from the 0 control to each condition, $i$, at each bin, $j$ (equation 2).

$$d_{j}\left( 0,i \right)=\sqrt{\sum_{k\in N} \left[ \ln\left( \frac{X_{ijk}}{g\left( X_{ij} \right)} \right)-\ln\left( \frac{X_{0jk}}{g\left( X_{0j} \right)} \right) \right]} (2)$$

$N=\left\{ A,C,G,T \right\}$ and $g(X_{ij})$ is the geometric mean for condition, $i$, and bin, $j$, across all four bases in $N$ (equation 3).

$$g\left( X_{ij} \right)=\sqrt[4]{\prod_{k\in N} X_{ijk}} (3)$$

For condition, $i$, and bin $j$, the signal, $s_{ij}$, is calculated as

$$s_{ij}=\frac{d_{j}\left( 0,i \right)-d_{j}\left( 0,0 \right)}{d_{j}\left( 0,1 \right)-d_{j}\left( 0,0 \right)}= \frac{d_{j}\left( 0,i \right)}{d_{j}\left( 0,1 \right)} (4)$$

where $d_{j}\left( 0,1 \right)$ is the Aitchison distance between the 0 control base composition and 1 control base composition at bin, $j$. $d_{j}\left( 0,0 \right)=0$ for all $j$. If there were multiple replicates for the 0 control, their average composition was used for $X_{0j}$ ($\mathrm{and}X_{0jk}$) in equation 2. If there were multiple replicates for the 1 control, their average composition was similarly used to calculate $d_{j}\left( 0,1 \right)$ in equation 4.

Next, the switch times were estimated for each condition, $i$, which contains a change in signal, $s_{ij}$, (e.g. via addition of Co halfway through the reaction). For experiments with more than one change (e.g. 0 🡪 1 🡪 0), a more sophisticated approach was used and is detailed below. However, the following simpler, more intuitive approach was used to predict switch times for 0 🡪 1 and 1 🡪 0.

Switch times were estimated for a given condition, $i$, by (1) finding $j_{i}^{*}$, the average location across all the sequences (bin position, $j$) at which half the 1 control signal is reached (i.e. $s_{ij}=0.5$), (2) calculating $\alpha$, the ratio of the average rate of nucleotide addition for the 0 and 1 controls, and (3) using $j_{i}^{*}$ and $\alpha$ to calculate the switch time, $t_{i}^{*}$, using equations 5 and 6. For a derivation of equation 5, see supplementary methods.

$$t_{i}^{*}=\frac{\alpha t_{expt}}{\frac{1}{j_{i}^{*}}+\alpha-1} (5)$$

where

$$\alpha=\frac{\bar{r_{a,ctrl}}}{\bar{r_{b,ctrl}}} (6)$$

$\bar{r_{a,ctrl}}$ is the average synthesis rate of the first environmental condition before the switch. For example, $\bar{r_{a,ctrl}}$ would be calculated using the 0 control for the condition, 0 🡪 1, but the 1 control for the condition, 1 🡪 0. The average synthesis rate is calculated by dividing the average extension length by the duration of the experiment. $\bar{r_{b,ctrl}}$ is the average synthesis rate for the second environmental condition (after the switch).

**Timepoint prediction for 0🡪1🡪0 multiple fluctuations experiment:**

To predict the Co^2+^ condition in the 0🡪1🡪0 experiment, we used a slight variation of the algorithm we developed in Glaser et al. for decoding continuous concentrations^4^. The input to this algorithm is the amount of signal on every nucleotide. Here, the signal is $s_{ij}$ from the previous section. The algorithm uses this information to predict continuous values of Co^2+^ between 0 and 1 for all time points that are most likely to produce the amount of signal on the nucleotides. When making these predictions, in the optimization cost function, we weighted each nucleotide added by the number of times a nucleotide occurred at that position (e.g. if there were 100000 instances of a nucleotide at position 1 and 5000 instances of a nucleotide being present at position 100, then the importance of their signals was weighted accordingly). To binarize the predictions, we set a threshold of 0.5. To be able to predict the values of Co^2+^, the algorithm requires knowledge of the expected amount of signal in the 0 and 1 control conditions. Here, this is the average signal across nucleotides in the 0 or 1 control experiments. The algorithm also requires knowledge of the rate of nucleotide addition. Here, we fit an inverse Gaussian distribution to the average experimental dNTP addition rate distribution (the distribution of the sequence lengths divided by the experiment time) from the control experiments. Note that this algorithm also assumes that the rate of dNTP addition is independent of the cation concentration. Thus, when making predictions in the 0🡪1🡪0 experiment, we do not account for differences in the rate of dNTP addition distributions between the 0 and 1 conditions. A future algorithm that takes this difference into account could yield more accurate predictions.

***In silico* simulations of experiments with multiple pulses:**

Using the average dNTP addition rate from experiments, and the amount of signal in the control conditions, we simulated additional experiments. Each simulated experiment had at least 3 pulses (instances of being in either the 1 or 0 condition), where each bit was randomly chosen to be 0 or 1. All nucleotides that were added during the 0 or 1 condition had the signal associated with these control conditions. More specifically, to account for the experimental variability in signals within a given control condition, nucleotide signals were sampled from a Normal distribution determined by the experimental variability of nucleotide signals within the control conditions. We calculated the variability in two ways, corresponding to the two curves in fig. S14. In one, the variability was calculated across the first 100 nucleotides, in which there were at least 2000 recordings of all base numbers. In the second, the variability was calculated across the first 50 nucleotides, in which there were at least 60000 recordings of all base numbers. Using the signal of the simulated nucleotides, we used the algorithm we developed in Glaser et al. for decoding binary concentrations^4^. The accuracy is the percentage of pulses correctly classified as 0 or 1.

Supplementary Text

**1. Extension reaction with individual dNTPs for testing effect of Co^2+^:**

For initial testing to show Co^2+^ dependent dNTP preference change the ssDNA substrate used was AMD006: 5’AGGCTAGTCGTCTGTATAGG3’. Total reaction volume was 25 μL with 0.1 µM ssDNA substrate, 1x NEB TdT reaction buffer, and 0.1 mM of each dNTP tested. Final concentration of CoCl_2_ in the test reaction was 0.25 mM. Reactions were initiated by addition of 5 units of TdT per reaction. Reactions were run for 30 minutes at 37 °C and stopped by boiling at 70 °C for 10 minutes. Then, 8 μL of the reaction was mixed with 12 μL of TBE-Urea loading dye and boiled for 10 minutes at 100 °C. All of the diluted extension reaction was then loaded onto 30 μL, 10 well 10% TBE-Urea Gel (Bio-Rad) and run for 40 minutes at 200 V. Immediately after the run was over, the gel was stained with Sybr Gold for 15 minutes and imaged on ImageQuant BioRad.

**2. Extension reactions for 1🡪0 set-up:**

Mg^2+^ only for 1 hour (signal 0) and Mg^2+^+Co^2+^ for 1 hour (signal 1) were set-up as regular extension reactions mentioned in Materials and Methods. The 1🡪0 reactions where the signal changed from 1 to 0 at 40 minutes were put through a ssDNA wash step at 40 minutes. ssDNA wash to remove cations, TdT and dNTPs was done exactly as mentioned in Materials and Methods. Reactions were all run for 1 hour with six replicates. Signal 0 and signal 1 controls were run for 1 hour for each set-up in triplicates and also put through the ssDNA wash step at 40 minutes.

**3. Derivation of Equation 5**

We start by deriving the equations for the average rate before the switch ($r_{A})$ and after the switch ($r_{B})$ for condition, $i$:

$$r_{a,i}=\frac{j_{i}^{*}}{t_{i}^{*}} (1a)$$

$$r_{b,i}=\frac{1-j_{i}^{*}}{t_{expt}-t_{i}^{*}} (2a)$$

where $j_{i}^{*}$ is the average location in the sequences (length fraction, 0 to 1) at which the signal, $s_{ij}$, reaches 0.5 (Equation 4), $t_{i}^{*}$ is the switch time, and $t_{expt}$ is the total duration of the experiment. Because we can estimate $r_{a,i}$ and $r_{b,i}$ from average rates of the 0 and 1 controls across replicates ($\bar{r_{a,ctrl}}$ and $\bar{r_{b,ctrl}}$), we can use their ratio to combine equation 1a and 2a, above to write

$$\frac{\bar{r_{a,ctrl}}}{\bar{r_{b,ctrl}}}\approx\frac{r_{a,i}}{r_{b,i}}=\frac{j_{i}^{*}}{t_{i}^{*}}\left( \frac{t_{expt}-t_{i}^{*}}{1-j_{i}^{*}} \right) (3a)$$

Solving for $t_{i}^{*}$, we get equation 5:

$$t_{i}^{*}=\frac{\alpha t_{expt}}{\frac{1}{j_{i}^{*}}+\alpha-1} (5)$$

where

$$\alpha=\frac{\bar{r_{b,ctrl}}}{\bar{r_{a,ctrl}}} (4a)$$

We use equation 5 for time prediction ($t_{i}^{*}$) after calculating $j_{i}^{*}$ for a given condition and $\alpha$ from the 0 and 1 controls. In equation 4a, $a$ is the first condition before the switch (0 or 1) and $b$ is the condition after the switch (1 or 0).

**4. Extensions reaction set-up for calculating rate of dNTP addition:**

Each extension reaction consisted of a final concentration of 10 μM initiating ssDNA substrate, 1 mM dNTP mix (each dNTP at 1 mM final concentration), 1x NEB TdT reaction buffer, and 10 units of TdT to a final volume of 50 μL. The ssDNA substrate used for this extension reaction was CS1_5N: 5’ACACTGACGACATGGTTCTACA(N1:25154515)(N1)(N1)(N1)(N1)3’. We have shown (fig. S9, A and C) that the identity of the last 5 bases on the 3’ end of the substrate affects the identity of the dNTP added to the ssDNA substrate. Thus, we purchased a ssDNA substrate (CS1_5N) with the last 5 bases having the base composition same as TdT dNTP preference under signal 0 (25% dATP, 15% dCTP, 45% dGTP and 15% dTTP). We happen to use this primer for this set-up, but we do not think the identity of the primer affects the rate of dNTP addition. The reactions were initiated upon addition of TdT and run at 37 °C for 2 hours. 2 μL of sample was collected and immediately frozen (on ice, 0 °C) at 30 s, 1 min, 2 min, 3 min, 4 min, 5 min, 10 min, 20 min, 30 min, 45 min, 60 min, 92 min and 120 min. Subsequently, each sample was put through the ligation and Illumina library generation process as mentioned in Materials and Methods.

**5. 0🡪1 at 10 minutes test set-up for checking ssDNA clean-up kit bias:**

Mg^2+^ only for 1 hour (signal 0) and Mg^2+^+Co^2+^ for 1 hour (signal 1) were set up as regular extension reactions mentioned in Materials and Methods. The 0🡪1 reactions where the signal changed from 0 to 1 during the 1 hour extension were run starting with 45 μL with Mg^2+^ only. 5 μL of 2.5 mM CoCl_2_ was added at 10 min. Reactions were all run for 1 hour in triplicates. Fresh signal 0 and signal 1 controls were run for 1 hour with each set-up. 2 μL of extension reaction was used directly for ligation (subsequently referred to as “No Wash” set of samples). Ligation and subsequent PCR steps for Illumina library generation were followed as mentioned in Materials and Methods. Rest of the 48 uL of extension reaction was washed using the ssDNA clean-up kit. Protocol was followed as mentioned in the kit. ssDNA was eluted into 25 μL of ddH_2_O and 2 μL of that was used for ligation (subsequently referred to as “Wash” set of samples). Ligation and subsequent PCR steps for Illumina library generation were followed as mentioned in Materials and Methods. Data obtained from Illumina sequencing was analyzed for the “No Wash” and “Wash” set of samples. Further, switch time calculations were carried out as mentioned previously (see fig. S13 for predicted switch times).

Supplementary Figures

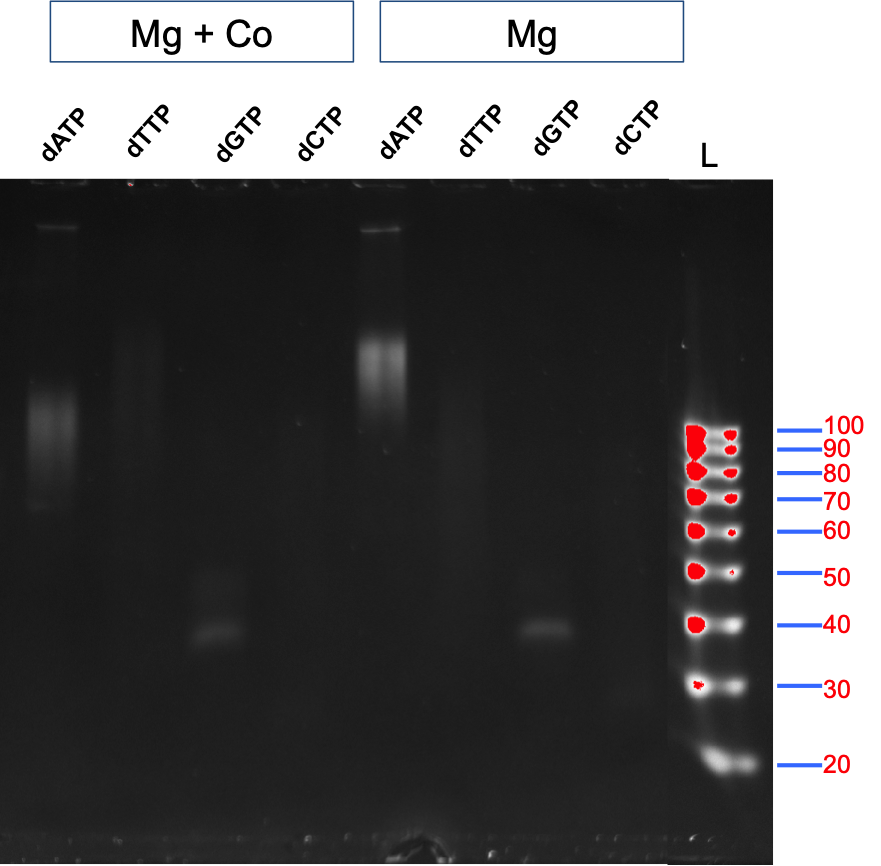

**Figure S1: Testing change in individual dNTP preference upon Co^2+^ addition**

ssDNA substrate extensions carried out by TdT using just dATP, dTTP, dGTP, or dCTP in presence of Mg^2+^+Co^2+^ (first 4 lanes) or in presence of just Mg^2+^ (next 4 lanes) were run on a gel. “L” is ssDNA size marker. Reactions were carried out as mentioned in supplementary text.

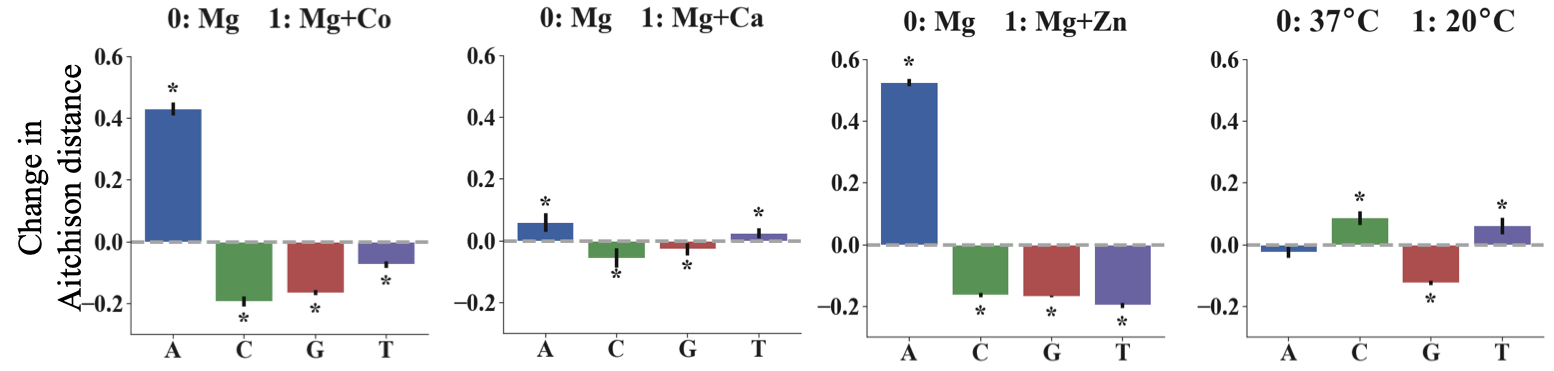

**Figure S2: Change in Aitchison distance for dATP, dCTP, dGTP and dTTP incorporation by TdT in the presence or absence of various signals.** Signal 0 is always 10 mM Mg^2+^ at 37 °C for 1 hour. Signal 1 was, going from left to right: (1) 10 mM Mg^2+^ + 0.25 mM Co^2+^ at 37 °C for 1 hour; (2) 10 mM Mg^2+^ + 1 mM Ca^2+^ at 37 °C for 1 hour; (3) 10 mM Mg + 20 μM Zn^2+^ at 37 °C for 1 hour; and (4) 10 mM Mg^2+^ at 20 °C for 1 hour. Error bars show two standard deviations of the mean Aitchison distance. Statistical significance was assessed after first transforming the data into Aitchison space which makes each dNTP frequency change statistically independent of the others. All base incorporation changes were found to be statistically significant, as shown by asterisks, (α = 0.01) except for the change in dATP for 20 °C signal (p-value = 0.019).

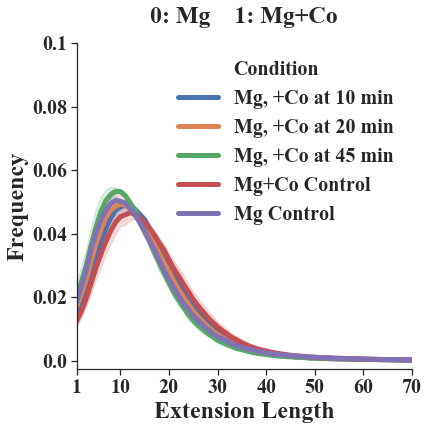

**Figure S3: Length distribution of extensions upon addition of Co^2+^ based on NGS data.**

We calculated the mean frequency distribution of extension lengths for each condition (three biological replicates for each condition). Addition of Co^2+^ did not change the length distribution significantly.

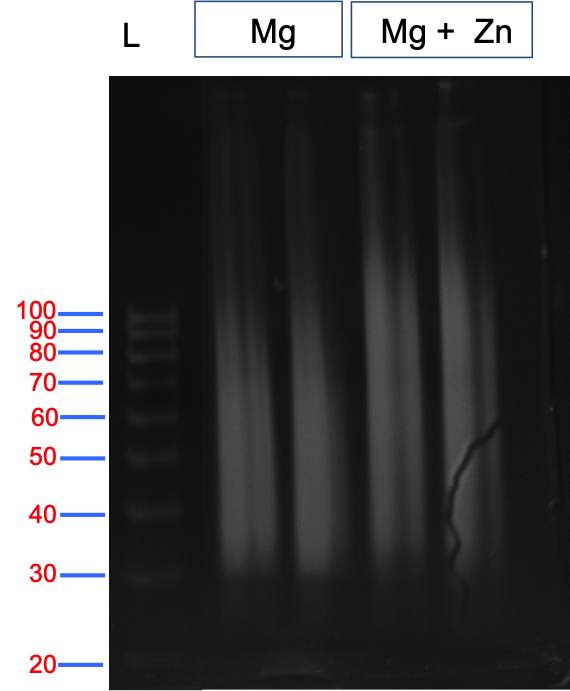

**Figure S4: Length distribution of extensions upon addition of Zn^2+^ as seen on ssDNA gel**

Extension reactions were run as mentioned in Materials and Methods section. Two biological replicates per test condition were then loaded onto a ssDNA gel (Mg^2+^ on left and Mg^2+^+Zn^2+^ on right). Addition of Zn^2+^ increases the overall lengths of the extensions.

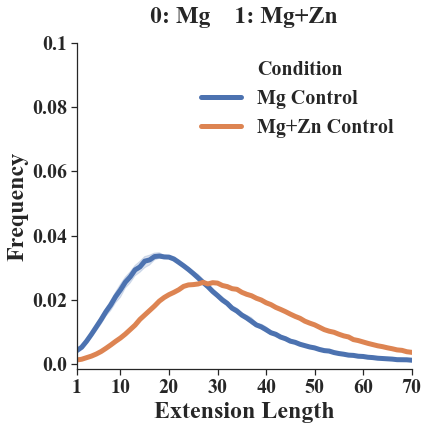

**Figure S5: Length distribution of extensions upon addition of Zn^2+^ based on NGS data**

We calculated the mean frequency distribution of extension lengths for each condition (three biological replicates for each condition). Addition of Zn^2+^ caused a shift in probability distribution toward longer lengths.

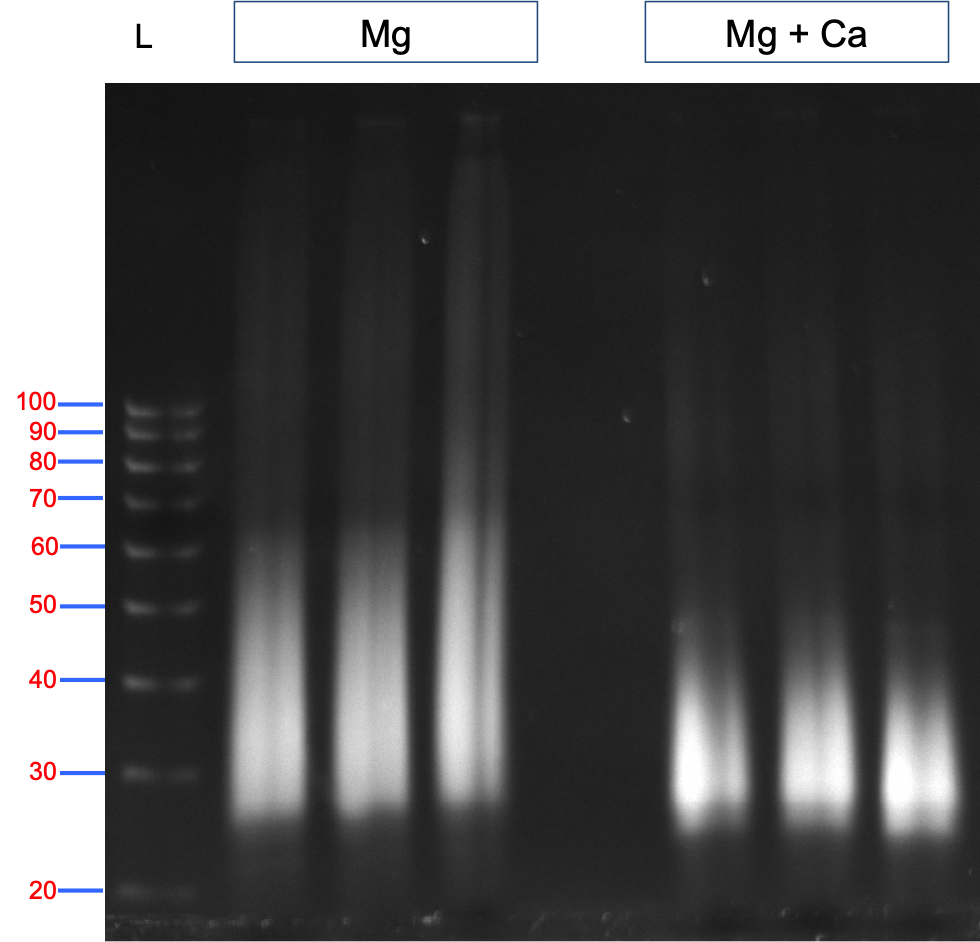

**Figure S6: Length distribution of extensions upon addition of Ca^2+^ as seen on ssDNA gel**

Extension reactions were run as mentioned in Materials and Methods section. Three biological replicates per test condition were then loaded on a ssDNA gel (Mg^2+^ on left and Mg^2+^+Ca^2+^ on right). Addition of Ca^2+^ decreases the overall lengths of the extensions.

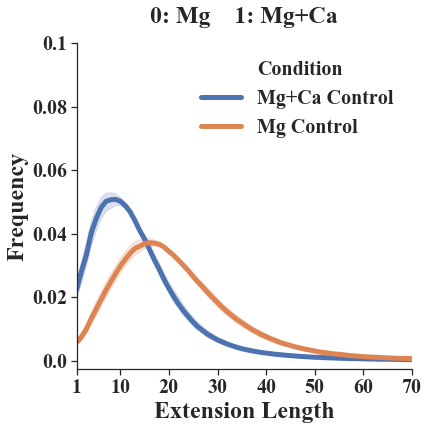

**Figure S7: Length distribution of extensions upon addition of Ca^2+^ based on NGS data.**

We calculated the mean frequency distribution of extension lengths for each condition (three biological replicates for each condition). Addition of Ca^2+^ caused a shift toward shorter lengths.

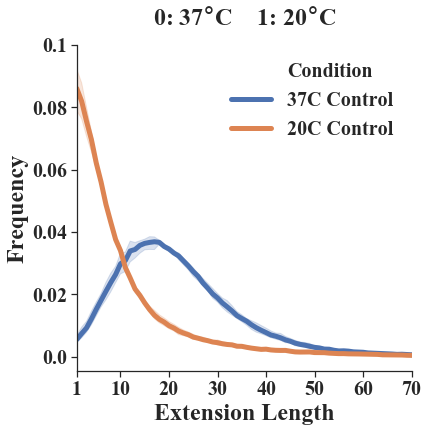

**Figure S8: Length distribution of extensions upon using temperature as a signal based on NGS data**

We calculated the mean frequency distribution of extension lengths for each condition (three biological replicates for each condition). Reducing the temperature of the extension reaction to 20 °C caused a shift toward shorter lengths.

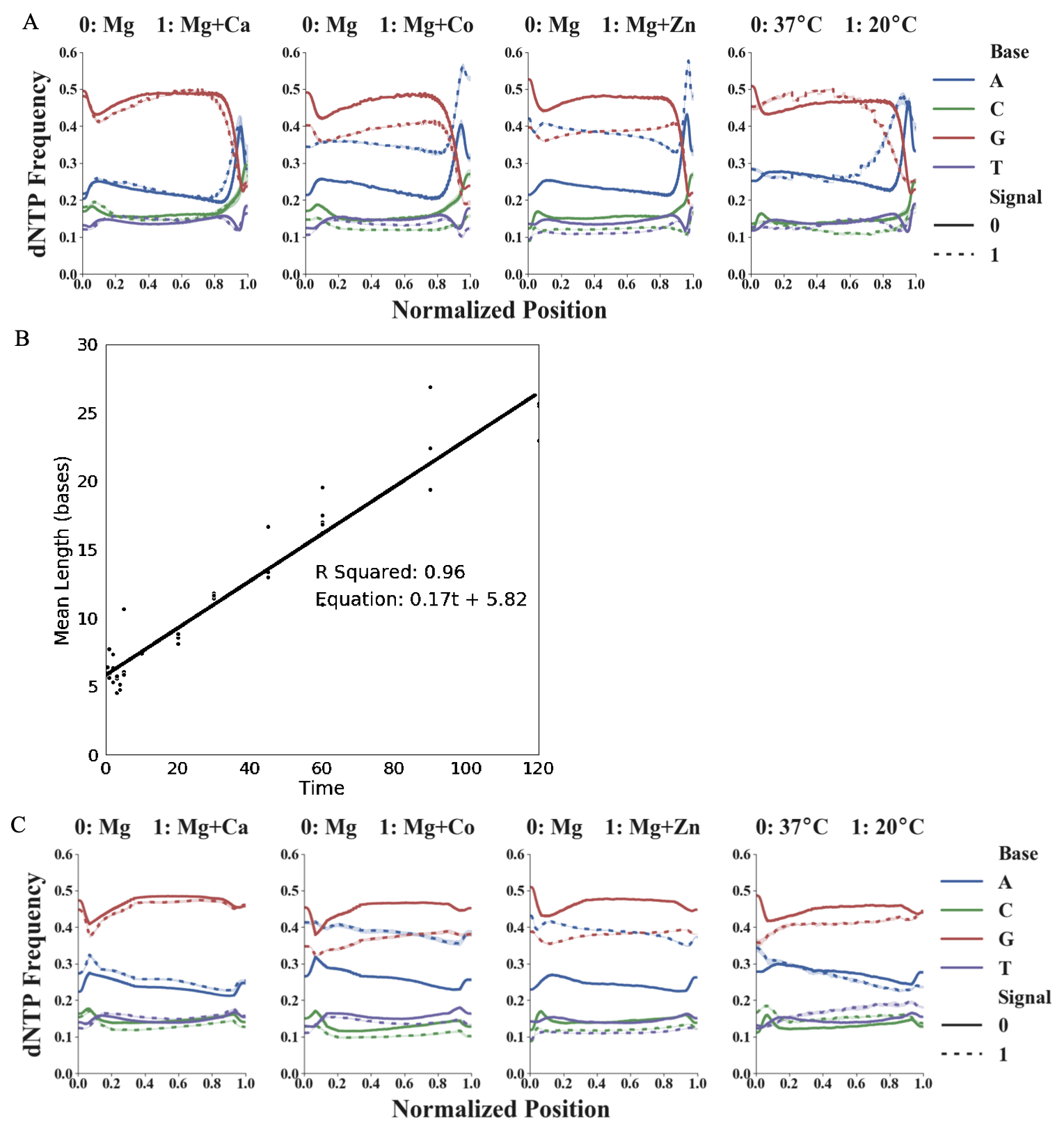

**Figure S9: Anomalous dNTP composition initially found at the end of reads and rate of reaction measured for extensions with only Mg^2+^ present**

We observed a significant change in the individual dNTP frequency towards the ends of the ssDNA sequences synthesized. (A) Presents the significant change observed near the end of all reads with all the signals tested. Since we directly use 2 µL of extension reaction for ligation, the diluted TdT seems to be adding dNTPs to the ssDNA after the recording experiment, during the 16-hour ligation step. (B) To prove that these dNTPs were not added during the extension reaction (i.e. after the reaction), we sampled extension reactions (with Mg^2+^ only) at several time points (Supplementary Text). We then calculated the mean extension length at each timepoint and applied a linear regression. The R^2^ value of 0.96 for a straight line indicates that our assumption of constant rate (assuming input signal does not change) is valid. The slope of 0.17 reveals an average incorporation rate of 0.17 dNTPs/minute for this condition. Most importantly, the intercept of 5.82 indicates addition of 5.82 dNTPs (on average) either before or after the extension reaction. These are almost certainly being added after the extension reaction during the ligation step, which we conclude based on the anomalous behavior we see at the end of sequences in Panel A. (C) We created plots of the data from Panel A after trimming off last few dNTPs. See Materials and Methods for details on how these 5.8 bases were trimmed from the end of all sequences before further analysis.

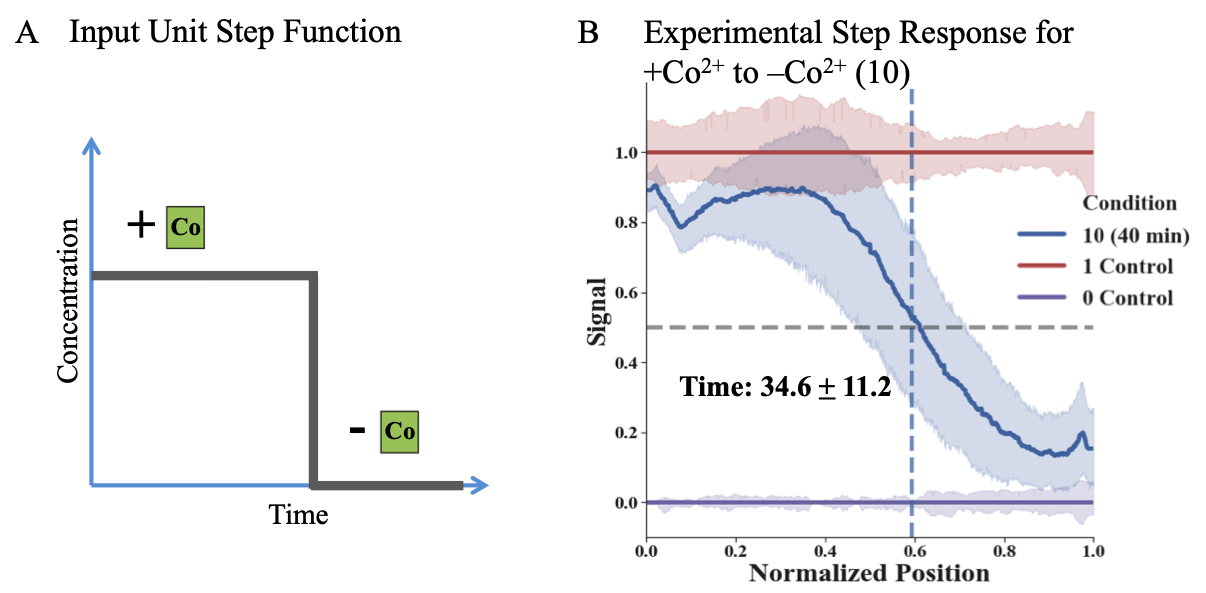

**Figure S10: Recording a single 1🡪0 step change in Co^2+^ concentration onto ssDNA *in vitro.***

(A) Representative input unit step function used in our experiments by changing concentration of Co^2+^ from 0.25 mM to 0 mM during a TdT-based DNA synthesis reaction while keeping Mg^2+^ concentration and reaction temperature constant. Co^2+^ was washed away at 40 minutes during a 60 minute extension reaction. (B) Experimental data for 1🡪0 step change. This plot shows there is a difference in the preference of dNTP incorporated by TdT in the Mg^2+^ (purple) and Mg^2+^+Co^2+^ (red) control conditions (where the signal (Co^2+^ is not added or removed throughout the extension reaction). The plot further shows the changes from 1🡪0 for Co^2+^ removed at 40 minutes (blue). We were able to get a switch time of 34.6 minutes with a std. dev. of 11.2 minutes using methods used for 0🡪1 switch time predictions (mean calculated across 6 biological replicates). We suspect the possible reason for higher variance in time prediction for this set-up was due to the ssDNA wash step at 40 minutes as discussed in fig. S13.

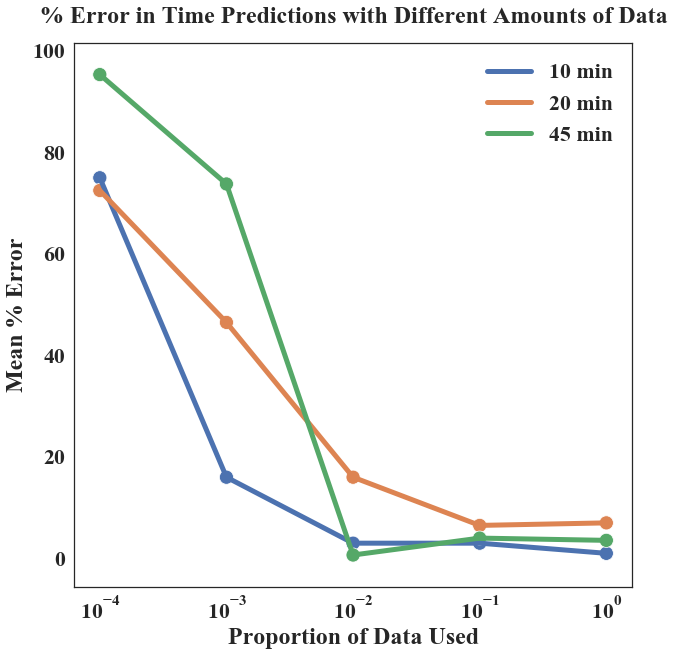

**Figure S11: Mean % error in time prediction for 0🡪1 (Mg^2+^ to Mg^2+^+Co^2+^) data when different proportions of experimental data are used for time prediction (data is randomly sampled)**

To get an estimate about how the accuracy of time prediction will vary with the number of DNA sequences analyzed we randomly sampled different proportions of experimental data obtained from the 0🡪1 setup. Roughly 600,000 sequences were sequenced for each reaction. Good prediction is obtained when at least 6,000 (1% of the original data) sequences are used for each reaction. The mean extension length was roughly 15 bases (in 60 minutes) for all conditions.

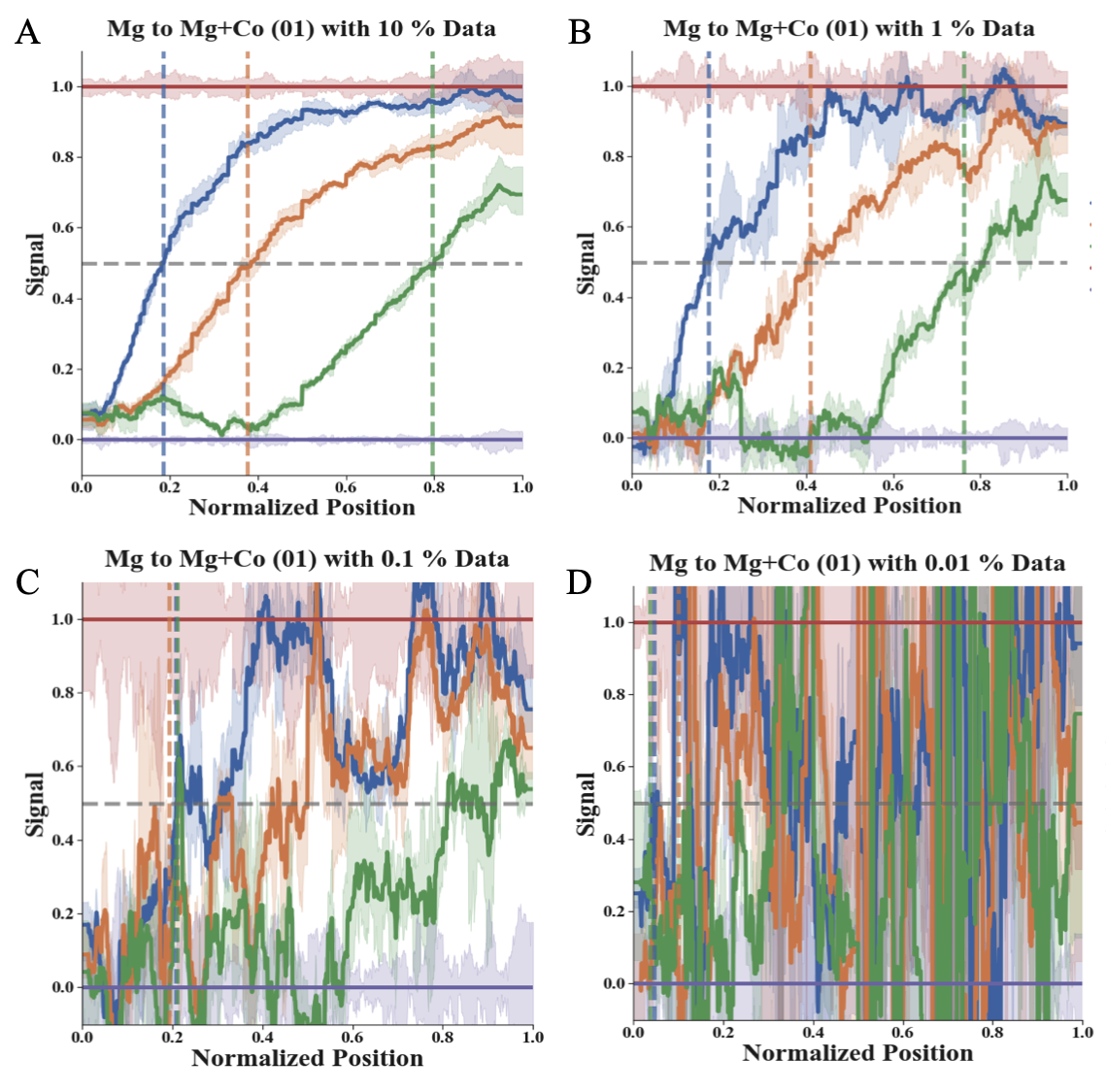

**Figure S12: Plots showing 0🡪1 data when different percentages of experimental data were randomly sampled**

(A) 10% of sequences (roughly 60,000 reads) obtained from the NGS data for the 0🡪1 set-up were plotted for calculating switch times. (B) 1% of sequences (roughly 6,000 reads) obtained from the NGS data set for the 0🡪1 set-up were plotted for calculating switch times. (C) 0.1% of sequences (roughly 600 reads) obtained from the NGS data set for the 0🡪1 set-up were plotted for calculating switch times. (D) 0.01% of sequences (roughly 60 reads) obtained from the NGS data set for the 0🡪1 set-up were plotted for calculating switch times. It is important to note that sequences were chosen randomly. For reference check Fig. 2C in main text, where 100% of the NGS data was plotted. Exact time predictions along with standard deviations can be found in table S1. fig. S11 shows error in time predictions for each panel. The mean extension length was roughly 15 bases.

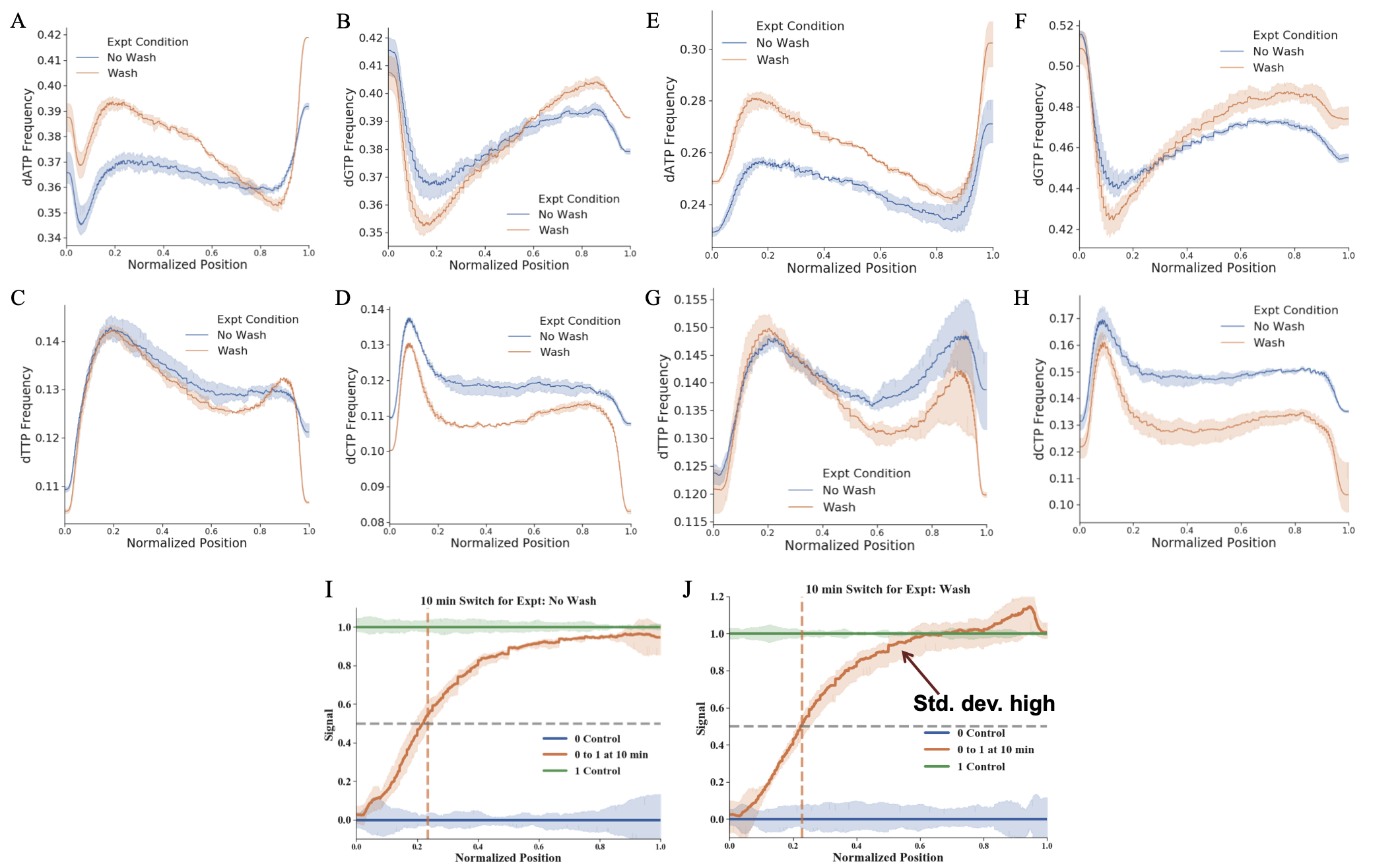

**Figure S13: dNTP bias & variablity introduced by ssDNA wash columns**

We present here a comparison of the composition of sequences retained when the extension reactions were directly used for ligation (“No Wash”) vs. when the same extensions were put through a ssDNA wash kit (“Wash”). (A), (B), (C) and (D) show individual plots of each nucleotide frequency seen in extension reactions between No Wash vs Wash conditions for just Mg^2+^ extensions. (E), (F), (G) and (H) show individual plots of each nucleotide frequency seen in extension reactions between No Wash vs Wash conditions for Mg^2+^+ Co^2+^ extensions. We observed a bias in overall dNTP content introduced by the columns used for ssDNA clean-up, when the reactions were washed after the recording experiment was stopped. ssDNA sequences with certain dNTP compositions were preferentially retained on the columns. (I) and (J) are plots for time prediction for No Wash and Wash condition respectively. We carried out an input signal of Co^2+^ 0🡪1 at 10 minutes for a 1 hour extension. We obtained a time prediction of 12.8 minutes with 1.8 min std. dev. for No Wash condition. We obtained a time prediction of 12.4 min with a std. dev. of 1.2 min for the Wash condition. While the time predictions were very similar, there is a clear increase in variability (std. dev.) for the later part of the signal recorded in (J) as compared to (I) (shown with a red arrow). Taken together, such biases and variability when introduced during the wash step for 0🡪1🡪0 experiment at 40 minutes for replacing +Co^2+^ buffers with –Co^2+^ buffers (See Materials and Methods: **Extension reactions for 0🡪1🡪0 set-up)** would cause more noise for the final 20 minutes of the recording.

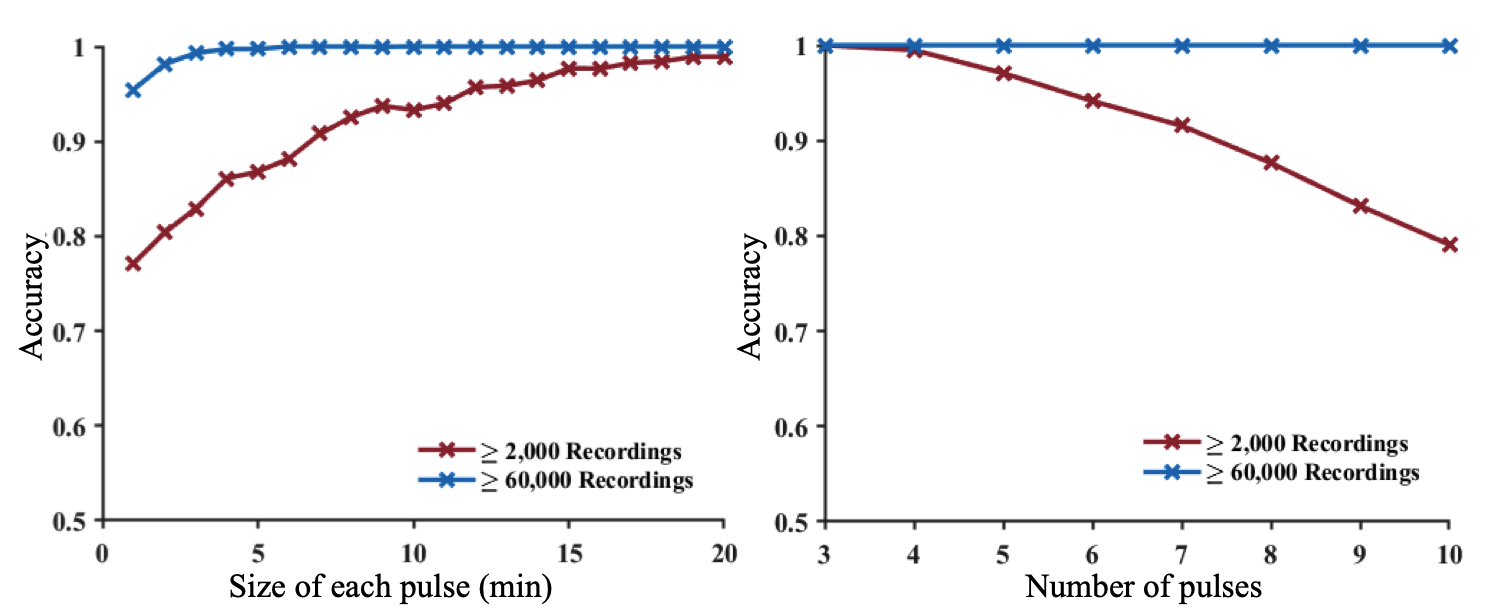

**Figure S14: *In silico* characterization of shortest resolvable pulses and highest number of consecutive pulses that could be resolved using TURTLES.**

*In silico* simulations based on experimental TdT parameters. For all simulations, we calculated the variability in two ways. The variability was either calculated across the first 100 nucleotides, in which there were at least 2000 recordings of all base numbers (red curve) or across the first 50 nucleotides, in which there were at least 60000 recordings of all base numbers (blue curve).(A) *In silico* characterization of shortest pulses resolvable. Simulations were carried out where the number of pulses (instances of being in the 0 or condition) was kept constant at 6 while the resolution per pulse was varied. Even in the high variability condition, 1 minute resolution is possible with > 75% accuracy, and 10 minutes resolution is possible with > 90% accuracy. (B) *In silico* characterization of number of consecutive pulses able to be resolved. Simulations were carried out with the duration of each pulse constant at 10 minutes while varying the number of total pulses. The accuracy of resolving this signal was calculated across simulations. Even in the high variability condition, 3 pulses can be resolved with almost 100% accuracy and 10 pulses of 10 minutes each can be resolved at about 80% accuracy.

**Supplementary Table**

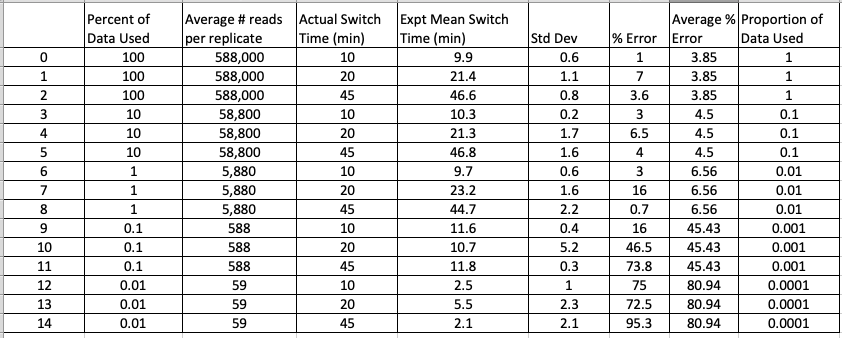

**Table S1: Table showing time predictions obtained for 0🡪1 data when different percentages of experimental data were randomly sampled**

To get an estimate about how the accuracy of time prediction will vary with the number of DNA sequences analyzed we randomly sampled different proportions of experimental data obtained from the 0🡪1 setup. Roughly 600,000 sequences were sequenced for each reaction. Good prediction is obtained when at least 6,000 (1% of the original data) sequences are used for each reaction with a standard deviation of about 1.4 minutes.
